# Draft genome of the Brazilian railroad worm *Phrixothrix hirtus* E.Olivier (Phengodidae: Coleoptera)

**DOI:** 10.1101/2021.12.01.470735

**Authors:** Danilo Trabuco Amaral, Yasuo Mitani, Isabel Aparecida Silva Bonatelli, Ricardo Cerri, Yoshihiro Ohmiya, Vadim Viviani

## Abstract

The Neotropical region is the richest in bioluminescent Coleoptera species, however, its bioluminescence megadiversity is still underexplored in terms of genomic organization and evolution, mainly within the Phengodidae family. The railroad worm *Phrixothrix hirtus* is an important biological model and symbolic species due to its bicolor bioluminescence, being the only organism that produces true red light among bioluminescent terrestrial species. Here, we performed the partial genome assembly of *P. hirtus*, combining short and long reads generated with Illumina sequencing, providing an important source of genomic information and a framework for comparative genomic analyses for the evaluation of the bioluminescent system in Elateroidea. The estimated genome size has ∼3.4Gb, 32% of GC content, and 67% of repetitive elements, being the largest genome described in the Elateroidea superfamily. Several events of gene family expansions associated with anatomical development and morphogenesis, as well as distinct *odorant-binding receptors* and *retrotransposable elements* were found in this genome. Similar molecular functions and biological processes are shared with other studied species of Elateriformia. Common genes putatively associated with bioluminescence production and control, including two luciferase genes that displayed 7 exons and 6 introns, and genes that could be involved in luciferin biosynthesis were found, indicating that there are no clear differences about the presence or absence of gene families associated with bioluminescence in Elateroidea. In *P. hirtus* the conversion of *L-* to *D-luciferin* seems to involve additional steps using a *Palmitoyl-CoA thioesterase* instead of an *Acyl-CoA synthetase*, which was found in Lampyridae species.

**Highlights:**

- First draft genome assembly of Phengodidae, the largest one described in Coleoptera;
- Gene family expansions associated with anatomical development and morphogenesis;
- Bioluminescent control and luciferin biosynthesis genes are common within Elateroidea;
- Despite similar bioluminescent system, metabolic routes may have evolved independently;

## 1. Introduction

The Neotropical region hosts the richest diversity of bioluminescent Coleoptera species in the world (Costa et al., 2010), including species of the three main families of Elateroidea: Elateridae, Lampyridae, and Phengodidae (Viviani, 2002; Rosa, 2010, Oba et al., 2011), and also two species of Staphylinidae (Costa et al., 2986; Rosa, 2010). Despite such richness, the Neotropical species are still poorly studied in terms of taxonomy, ecology, and evolution. Among the three main families, the less studied one is Phengodidae, which has 35 genera and 200 species distributed in three tribes, Pennicilloporini, Phengodini and Mastinocerini, the latter being found predominantly in the Neotropical region (Wittmer, 1976; O’Keefe, 2002; Viviani, 2002). The *Phrixothrix hirtus* E.Olivier (1909) railroad worm (Fig. 1a) is certainly the most spectacular example of bioluminescence in Phengodidae, with its unique red-light emitting cephalic and post-cephalic lanterns and yellow-green emitting lanterns in the body, having been used as an important biological model for biochemical and molecular studies of luciferases (Viviani et al., 1999a; 2001; 2004; 2013; 2021; Viviani and Ohmiya, 2000; Amaral et al., 2016; Bevilaqua et al., 2019). The bicolor bioluminescence is caused by the presence of luciferase isozymes in the head and lateral body lanterns (Viviani and Bechara, 1995), being an example of paralogy caused by events of gene duplication in Mastinocerini (Arnoldi et al., 2010).

**Figure 1.**
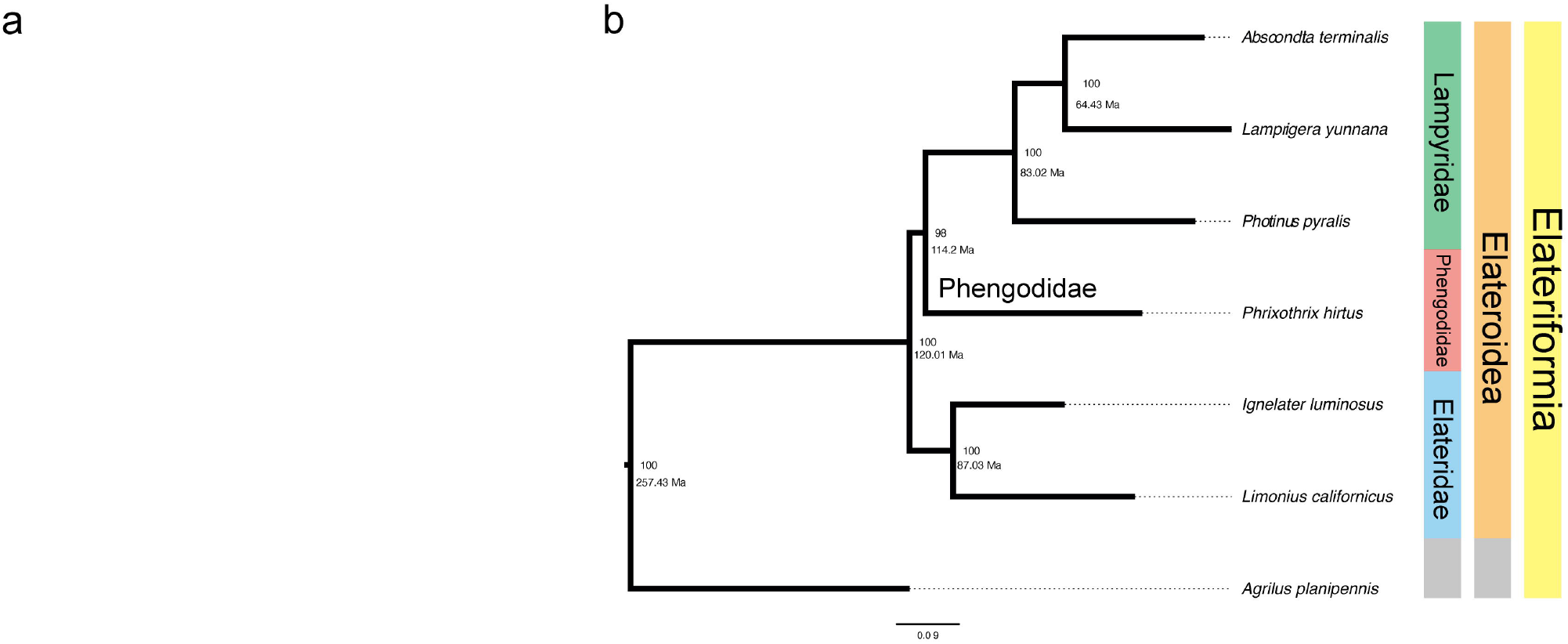
Phylogenetic context of Elateriformia. **a**. *Phrixothrix hirtus* railroad worm. **b** Phylogenetic tree depicting the relationship within Elateroidea bioluminescent species, with *Agrilus planipennis* (Buprestidae) as outgroup.

Genome analysis has been used to understand the evolutionary processes that drive the diversification and the evolution of the bioluminescence in Elateroidea (Braham and Wenzel, 2001; Day et al., 2004; Amaral et al., 2019a; 2019b). The genome size estimation in Elateroidea based on flow cytometry was also performed for all bioluminescent families (Hanrahan and Johnston, 2011; Liu et al., 2017; Lower et al., 2017). These studies suggested that the genome sizes among species of this superfamily range from 0.4 Gb to 2.2 Gb, and identified a positive relationship between the genome size and the amount of repetitive DNA. Recently, the genome sequence and assembly of *Photinus pyralis* firefly (Lampyridae) and *Ignelater luminosus* click-beetle (Elateridae) (Fallon et al., 2017; Fu et al., 2017; Andrews et al., 2020) were conducted and deposited in public genetic database repositories. These genomic analyses showed the divergence of the ancestral luciferase, supporting the independent origins of bioluminescence in Elateridae and Lampyridae. They also showed the existence of two distinct luciferase genes in *P. pyralis* firefly located in two chromosomes, suggesting events of gene duplication associated with a translocation between the chromosomes, which corroborates the presence of two luciferases isozymes in the lanterns and fat body in distinct life stages of fireflies (Strause and DeLuca, 1981; Viviani et al., 2008; Oba et al., 2010; Bessho-Uehara and Oba, 2017; Carvalho et al., 2020). In addition, genomic and transcriptomic analyses of two Palearctic firefly species, *Abscondita terminalis* and *Lamprigera yunnana*, indicated putative luciferin biosynthesis pathways in fireflies (Zhang et al., 2020), involving several gene products, which were also described in RNA-Seq analyses in other Elateroidea species (Vongsangnak et al., 2016; Amaral et al., 2017a,b;2018). Such studies brought important contributions and insights about the genome organization within Elateroidea, however, the functional genomic information remains limited to Lampyridae and Elateridae.

However, with the exception of RNA-Seq analysis of the lateral lanterns and fat body of *P. hirtus* which identified several luciferase-like enzymes in these tissues and a luciferase enzyme in photogenic tissues (Amaral et al., 2017a; 2017b; 2019b), no genome sequence analyses was conducted for the Phengodidae family yet. Genome sequence analysis in this family may improve the knowledge about the origin and evolution of the family, as well as the origin and control of bioluminescence.

Here we report the first draft genome assembly of a Phengodidae species, the South-American *Phrixothrix hirtus* E.Olivier railroad worm, using both genomic short-read and mate-pair libraries. This species displays the largest genome among Elateriformia species studied so far, as well as the presence of several transposable element families. With this draft genome, we produced a partial genome assembly, which is a novel important source of information for future structural, genetic, evolutionary studies, and for comparative genomic analyses in Coleoptera species evolution of bioluminescence in Elateroidea.

## 2. Results and Discussion

### 2.1. *de novo* Genome Assembly and Annotation

The genome of *P. hirtus* (Fig. 1a) is the seventh sequenced and available genome within the Elateroidea superfamily and the first one in the Phengodidae family. The phylogenetic analysis showed a strict relationship between Lampyridae and Phengodidae (Fig. 1b). Based on previous findings, Phengodidae diverged from the sister Palearctic family, Rhagophthalmidae, around 73.4 Ma, and from Lampyridae around 97.3 Ma (Amaral et al., 2019). The ultrametric tree performed by the r8s software suggested that the divergence between Phengodidae and Lampyridae occurred even earlier, around 114 Ma.

Here, we generated and assembled ∼190 Gb (56.8-fold coverage) from Illumina short reads and ∼65Gb (20.2-fold coverage) from Illumina mate-pair reads (Table S1 and S2). The assembly length, ∼3.40□Gb, was consistent with the *k*-mer estimate genome size (□3.4□Gb in Jellyfish2.0 and ∼3.35Gb in GenomeScope; Fig. S1). So far, this is the largest described genome among Elateroidea species (from 0.42Gb to 2.5Gb) and also in Coleoptera (from 0.15Gb to 2.7Gb) (Hanrahan and Johnston, 2011). Previously, we showed that the mtDNA genome of *P. hirtus* is also larger than those from other Elateroidea species, with duplication events and a larger control region (Amaral et al., 2016). Several hypotheses were suggested and tested to evaluate the genome size correlation in Coleoptera, including body size (morphological; Palmer and Petitpierre, 1996), chromosome number (Petitpierre et al., 1993), methylation rate (Lechner et al., 2013), and reproductive fitness (Arnqvist et al., 2015). However, the genome size correlation in railroad worms needs to be better explored, including new species and morphological, environmental, and genomic data.

The *P. hirtus* scaffolded genome is detailed in table S2. The gene prediction using Augustus resulted in 253,925 gene models in the first round of analysis, and 92.234 gene products in the second round of analysis. The predicted proteins were found against the SwissProt database, in which 87,967 genes were assigned to the putative function (Fig. S2). These observed GO terms are common in most arthropods, being responsible for basic physiologic processes and metabolic activities that are essential in Insecta. We did not identify any specialized function occurring only in *P. hirtus*. The genome contains approximately 70% complete single-copy orthologs and multi-copy orthologs, which indicate that most parts of genes were recovered. The percentage of observed GC content was ∼32% and the estimated heterozygosity was about 0.25% (Fig. S1). We obtained a total of ∼12 million elements of repetitive DNA (2,326,217,477 bp, or 2.4Gb), representing approximately 67% of the total length of the assembled scaffolds (Table S3).

We also completely assembled the circular mitogenome of *P. hirtus*, which displayed 20,303 bp, similarly to the previously studied in this species (Amaral et al., 2016), again being the largest Elateroidea mtDNA described. The mtDNA presented 2 ribosomal RNAs (rnaS and rnaL), 13 protein coding-genes (PCG), and 21 tRNAs (absence of tRNA-A; Table S4). A larger *A+T rich-region* (5,661 bp), which includes four partial copies of the *ND2* gene and the tRNA-Q was observed. Similar to the nuclear genome, these results also support the evidence of high dynamism of the *P. hirtus* genomes.

### 2.2. Repetitive element DNA content

The inter-species genome size variation could be the result of several gene/genome duplication and/or deletion events (Blommaert, 2020), however, the rearrangements and duplication of repetitive DNA sequences may also imply large genomes sizes (Talla et al., 2017). Studies are showing the role of these elements as potential substrates for new genes and their association with gene expression (e.g. epigenetic regulation), as well as a stress response (regulatory sequence) (Rech et al., 2019; Choi and Lee, 2020; Fedoroff, 2020). Thus, these elements could be major drivers of genome evolution in Eukaryotes (Quesneville, 2020).

According to BUSCO, there was no excess of duplicated genes (7.42% of duplicates). Nevertheless, the analysis of the landscape of repetitive and transposable elements (TE; Table S3) of *P. hirtus* showed that its genome has a great part of base-pair length composed by repetitive elements (∼67%; 2.4Gb). The percentages of TE content in Elateroidea vary from 19.8% (∼190Mb) in *A. lateralis* to 42.6% (∼180Mb) in *P. pyralis* (Fallon et al., 2018). In *P. hirtus*, the majority of these TE in *P. hirtus* were unclassified (46.83%) followed by the DNA transposons (12.3%). Recent studies have also discussed the variation of TE contents among Coleoptera, Diptera (6% in the *Belgica antarctica* to 58% in *Anopheles gambiae*), Hymenoptera (less than 6% in *Apis mellifera* and *Athalia rosae*), and Orthoptera (58% in *Locusta migratoria*), which suggested that the repetitive elements are highly dynamic in Insecta (Hazzouri et al., 2020).

### 2.3. Orthologous Analysis and Evolution of gene families in Elateriformia

The comparative genomic analysis evaluates the ortholog gene families represented in the genome to improve the understanding of the evolution within this superfamily. Here, the orthologous analysis using the amino acid sequences of Elateriformia species (*Abscondita terminalis, Lamprigera yunnana, Photinus pyralis* (Lampyridae), *Ignelater luminosus, Limonius californicus* (Elateridae), and *Agrilus planipennis* (Buprestidae)) against *P. hirtus* showed 17,862 ortholog gene families. Among them, 2,001 orthogroups (11.2% of the total) were commonly shared among all species (Fig. 2), with only 359 orthogroups displaying a single gene copy, which were applied to phylogenetic reconstruction (Fig. 1b); a total of 523 orthogroups in *P. hirtus* did not share direct orthology with the other six protein species. The bioluminescent species used in this study shared 2,477 orthogroups, however, only 440 displayed single-copy genes.

**Figure 2.**
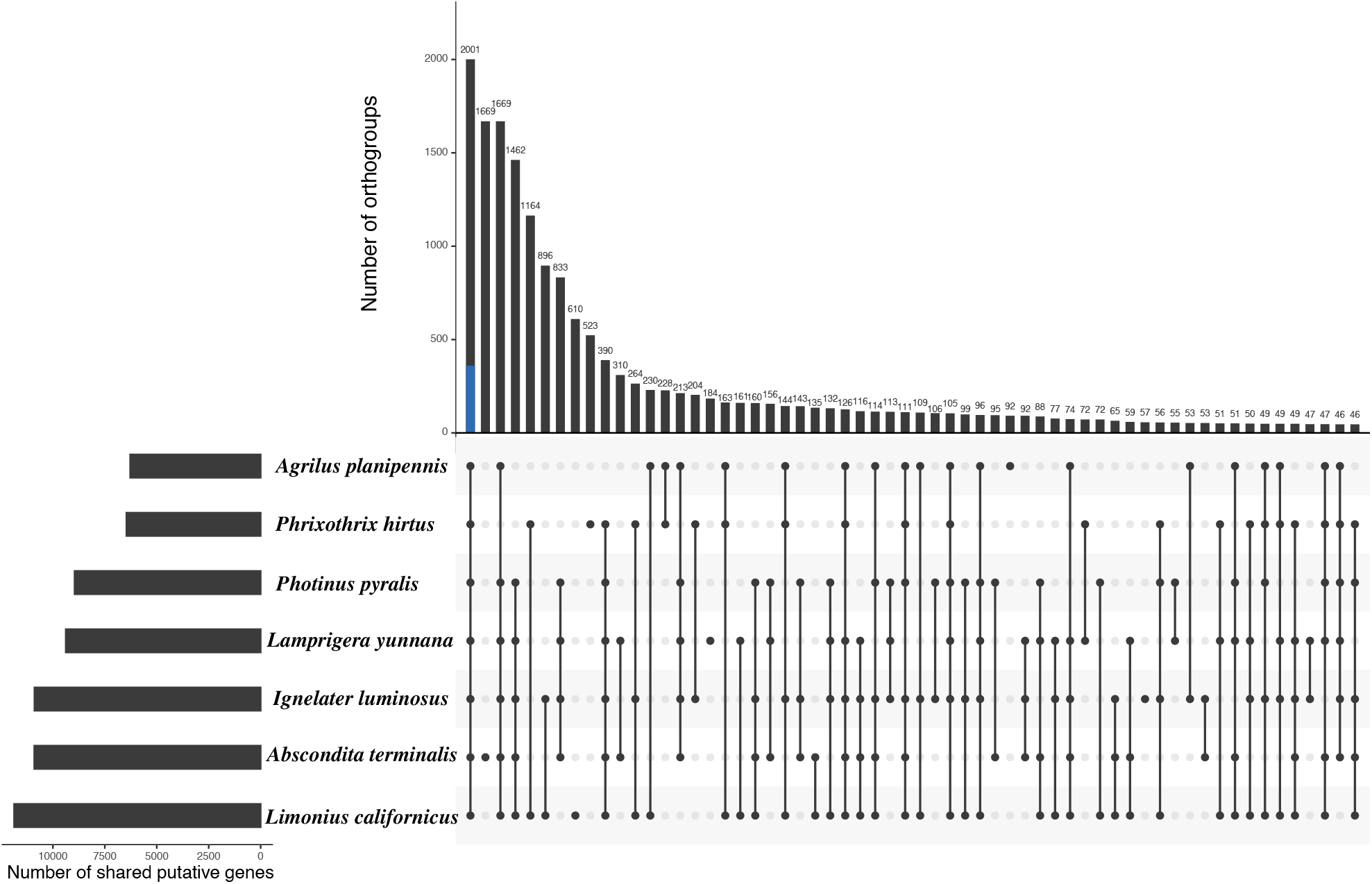
Barplot of shared and unique orthologues among Elateriformia genomes. In blue, the number of single-copy genes shared among the species in comparison to multiple copies orthogroups (in black).

The molecular function ontology of the predicted gene products shared among all Elateriformia species shows enrichment of catalytic activity and binding enzymes (Fig. S3A and S3B). Among catalytic activity, the hydrolase and transferase were highlighted, while among binding functions, ion-binding, heterocyclic binding, organic cyclic compound binding, and protein binding were highlighted (Table S5). Biological processes of the shared GO were mostly enriched for cellular processes (regulation of cellular process and cellular metabolic process), biological regulation (regulation of biological quality), regulation of biological process (regulation of metabolic process), and metabolic process (nitrogen compound metabolic process, organic substance metabolic process, and primary metabolic process) (Fig. S3A and S3C).

We separately evaluated the orthogroups associated only with the bioluminescent species of the Elateroidea (Table S6). Considering these five species, we obtained 2,477 orthogroups, whereas 476 were exclusively for bioluminescent individuals. We also annotated their gene ontology (Fig. S4). Here, we identified several families of general odorant-binding proteins (OBP), responsible for recognizing and transporting hydrophobic odorants to the antennal sensilla and activating the olfactory signal transduction pathway (Li et al., 2016). For many flashing firefly species such as *Photinus pyralis*, it is expected and suggested that the luminescence emission pattern (flash, continuum/glow, etc.) is the main factor responsible for intraspecific communication. However, in *Phrixothrix* railroad worms, as well as other Phengodidae, it is known that pheromone detection plays a major role in sexual attraction (Jacobson, 2012). The presence of these several OBP gene families supports that both the pheromones and bioluminescence are involved in the communication between Elateroidea species.

The comparison analysis among Elateriformia and Elateroidea did not show clear differences in the presence or absence of gene families among bioluminescent species and their closely related non-bioluminescent species, with the exception of OBP gene families in Phengodidae. These results suggest that both bioluminescent and non-bioluminescent species display similar groups of gene families, including luciferase and/or luciferase-like enzymes, genes associated with luciferin biosynthetic pathway (Niwa et al., 2006; Oba et al., 2013; Amaral et al., 2017a; 2017b; 2019b; Zhang et al, 2020), sulfotransferase (Fallon et al., 2016), etc. However, the transcription control and expression level of these genes could be the main determinants of the spatial and temporal control of the bioluminescence in Elateroidea, rather than a genomic feature.

### 2.4. Analysis of unique *P. hirtus* orthogroups

We observed 523 orthogroups in *P. hirtus* that did not share direct orthology with the other studied species. In the annotation of these gene families, however, only a small part of the orthogroups was assigned (∼12%) (Table S7). From them, several retrotransposable elements, such as LINE, MOS1, ATP Translocases, and PiggyBac, were present among these orthogroups. The high amount of these elements exclusively in the *P. hirtus* genome may explain the largest genome. The eukaryotic genome is highly dynamic, including events of gene duplication or even whole-genome duplication (Ting et al., 2004; Van de Peer et al., 2009; Mendivil and Ferrier, 2012). In arthropods, these mechanisms may have important evolutionary significance when associated with adaptation to environmental changes and the processes of biological and cellular regulation and simultaneous response to external stimulus (Kidwell, 2002; Chénais et al., 2012).

We also observed in *P. hirtus*, the gene families of *Craniofacial development protein* and *Heat Shock*, which seem to work in consonance to the development of wing dimorphism in arthropods (Carrol, 1995; Baral et al., 2019; Chen et al., 2019). In Phengodidae, including *P. hirtus* railroad worms, the sexual dimorphism is accentuated, with a clear distinction between male and female body shape and bioluminescent organs (Wing et al., 1984). The females are neotenic and paedomorphic (retention of larval traits), while the males undergo metamorphosis and develop wings in the adult stage. Furthermore, the males are much smaller than the females, besides the presence of wings. However, more molecular biochemical studies are necessary to understand the genes involved in metamorphosis and sexual differentiation in the family Phengodidae.

### 2.5. Expansion of ortholog gene families

In the past few years, comparative genomic studies showed the dynamic aspect of genome size and gene families in rapidly evolving groups of species, such as plants and insects (Zhang et al., 2018; Freitas and Neri, 2020; Hazzouri et al, 2020; Jiao, et al., 2020; Wang et al., 2020). The expansion and contraction of gene families seem to be pervasive and provide evidence that the copy number changes are associated with the natural selection acting under the particular adaptation of the species, such as changes in protein-coding and regulatory regions (Hahn et al., 2009). These gene families with complex gene duplication histories in lineages deserve great attention. Thus, we estimated the expansion and contraction of gene families among Elateriformia species.

The protein orthogroups obtained here were managed to identify signatures of expansion in gene families among Elateriformia. The number of rapidly evolving, expansions, and contractions gene families was displayed in Fig. 3. There are a total of 2,134 expanded and 7,581 contracted gene families among Elateroidea families internal branches. The largest number of contraction gene families was found in *P. hirtus* (9,783), while the largest number of expansion was found in *Limonius* (4,268). The branches with the largest numbers of rapidly evolving gene families are both leading to Elateridae and Lampyridae families (194 and 173, respectively). The ancestor branch of Elateroidea displayed 1,514 and 547 gene families with contractions and expansions, respectively. Details about the molecular function and biological processes of the gene families that displayed expansion within the Elateroidea superfamily are shown below.

**Figure 3.**
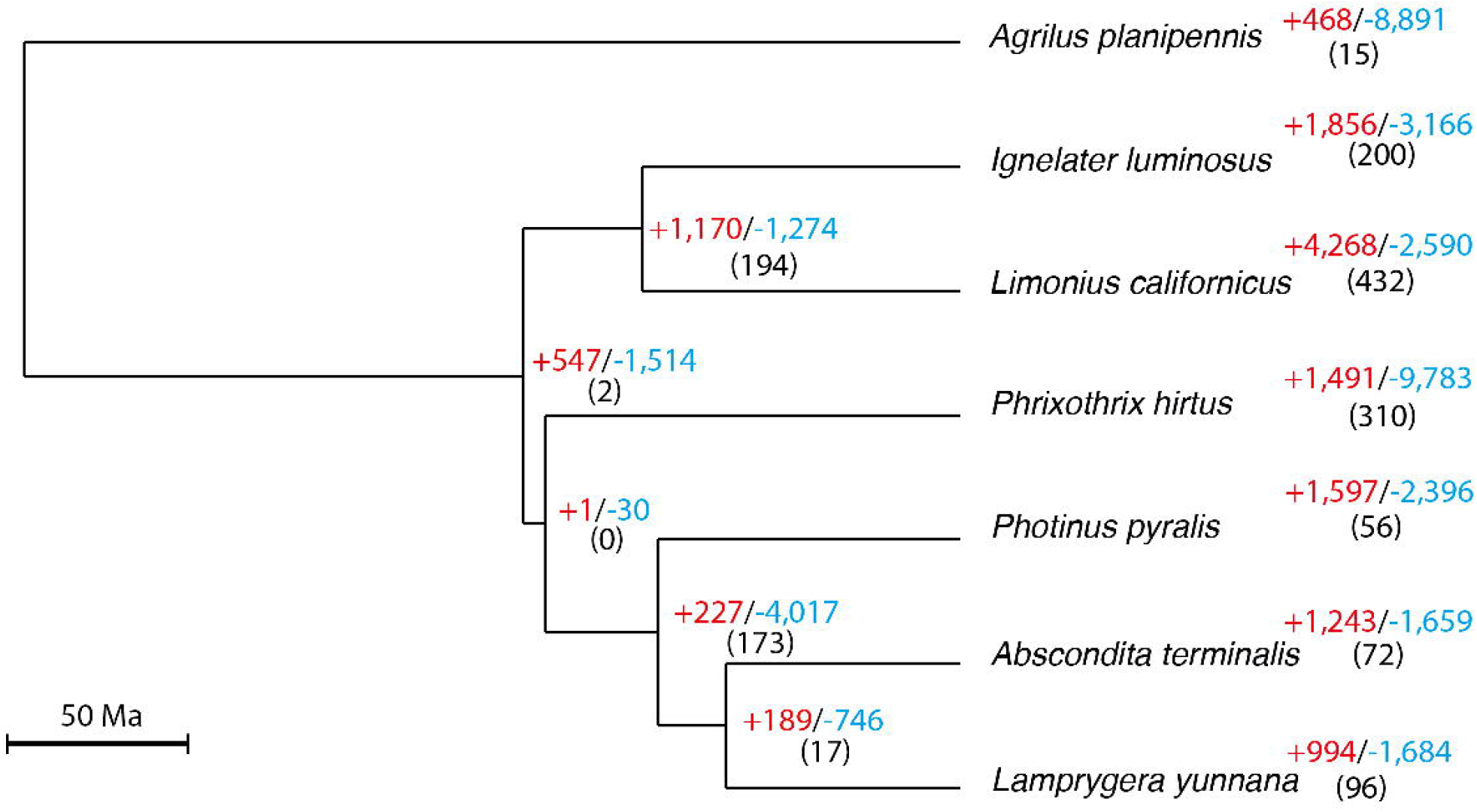
Summary tree shows the inference of the gene family evolution based on the expansion (+ blue) and contraction (-red) gene families in Elateriformia. The number between parentheses represents the number of rapidly evolving gene families.

#### Elateroidea

In the main branch of Elateroidea, we observed 547 expanded gene families. From them, a total of 465 gene families were annotated and classified concerning their molecular function (Table S8). The gene families are involved with catalytic activity (hydrolase and oxidoreductase activity) and binding (ion, organic cyclic compound, and heterocyclic compound binding), as well as metabolic processes (organic substance, nitrogen, and primary metabolic process), regulatory processes, and response to stimuli (Fig S5A). We found a high amount of hydrolases (several *lipase* members, *serine hydrolase, glutathione hydrolase, Acyl-CoA thioesterase*), transferases (*glutathione S-transferase, UDP-glucosyltransferase, Acyl-coenzyme A:cholesterol acyltransferase 2, Sulfotransferase, odorant binding)*, ligase (*E3 ubiquitin ligase*), and oxidoreductases (several *Cytochrome P450* members, *Fatty acyl-CoA reductase, luciferase, phenoloxidase*). Several of these gene families in the expansion are associated with detoxification and metabolism of xenobiotics (*cytochrome P450s, glutathione S-transferases, UDP-glucuronosyltransferases*, etc.) (Ahn et al., 2012; Zhu et al., 2016; Rane et al; 2019), and were widely identified among the transcriptome dataset of Elateroidea (Amaral et al., 2017a; 2017b; 2019b). These genes play fundamental roles in xenobiotic detoxification and degradation of distinct molecules related to an insect diet, detoxification of metabolic compounds, resistance to pesticides, degradation of hormones (Xue et al., 2020), and the degradation of the lucibufagins (defensive steroids), present in some fireflies species and used for defensive purposes (McKinley and Lower, 2020).

#### Elateridae

In the Elateridae branch, we identified and annotated 819 gene families (from 1,170; Table S9), which recovered quite similar patterns observed to the Elateroidea branch, with the high number of gene families related to the catalytic activity (hydrolase activity) and biding (ion, organic cyclic compound, and heterocyclic compound binding) and metabolic processes (organic substance, nitrogen, and primary metabolic process). We also observed a high amount of cellular processes (regulation and metabolic process; Fig S5B). Gene families such as *lipase* members, *serine hydrolase, glutathione S-transferase, UDP-glucosyltransferase, Sulfotransferase, luciferase, E3 ubiquitin ligase*, several *Cytochrome P450* members, and *Fatty acyl-CoA reductase* were the most common gene identified.However, we also observed the expansion of *phenoloxidase 1* gene family, *succinate CoA-ligase* (ADP-formin superfamily), and *craniofacial development protein 2* (relevant to wing-shed occurrence; Chen et al., 2019).

#### Agrypininae/Pyrophorini (bioluminescent click beetles *Ignelater luminosus*)

We evaluated the gene families’ expansion within the bioluminescent species *Ignelater luminosus* of Elateridae and obtained 1,856 gene families. A total of 1,366 were annotated (Table S10). The GO molecular function and biological processes were quite similar to the Elateridae branch (Fig S5C). Several *ubiquitin protein ligases, sulfotransferase* (five distinct gene families), *Cytochrome P450, UDP-glucosyltransferase*, and *glutathione S-transferase* were identified. Evaluating all other branches, the expansion of the luciferase gene family is present only in Elateridae, with the presence of several luciferases and luciferase-like enzymes (Fallon et al., 2018). The high amount of replication and transcription factors, *perithophin* (insect pathogenic protection), and *keratin-associated protein* (also observed in Lampyridae) were singular to the *I. luminosus*.

#### Denticollinae (Non-luminescent Elateridae *L. californicus*)

The non-luminescent click beetle *Limonius* displayed the highest number of expanded gene families among the studied species (4,268 with 2,146 annotated; Table S11). These gene families are associated with the binding molecular function (mainly ion binding) and regulation, metabolic, and stimulus of cellular processes (Fig S5D). Several families displayed more than 2 or 3 distinct gene families. The interesting finding is the expansion of two distinct families of *luciferase*, however, we did not identify the expansion of luciferase-like gene family, such as *4-coumarate-CoA ligase* and *succinate CoA-ligase* (ADP-formin superfamily), which were present in other genomes and also observed in recent transcriptome analysis luminescent and non-luminescent Elateridae species (Amaral et al., 2019b).

#### Lampyridae

A total of 256 (from 416) gene families were annotated (Table S12). Major molecular function and biological processes of gene families were catalytic activity, binding, and cellular processes (Fig S5E). We observed the expansion of *glutathione S-transferase, sulfotransferase* (three distinct gene families), *acyl-CoA thioesterase*, and *Cytochrome P450*, which are highlighted in other Elateroidea species. Here, we did not identify the expansion of the luciferase gene family, however, we observed the expansion of the *4-coumarate-CoA ligase*, which also belongs to the adenylate-forming family. This group of gene families was also observed in the transcriptome analysis of the three bioluminescent species of Elateroidea, Elateridae, Lampyridae, and Phengodidae (Amaral et al., 2017a, 2017b, 2019b). We identified the expansions of the *superoxide dismutase* (SOD), *cystathionine-B-synthase*, and *cysteine-rich protein 2-binding*. The SOD enzyme seems to be associated with the protection of lanterns during hyperoxia and oxidative stress and is recovered in high abundance in lanterns of firefly species (Barros and Bechara, 1998; Amaral et al., 2017b). The other two gene families are associated with cysteine availability, which is a precursor of luciferin biosynthesis in Elateroidea (Viviani et al., 2013; Kanie et al., 2016; Amaral et al., 2017a, 2017b, 2019b; Zhang et al., 2020).

#### Phengodidae (*Phrixothrix hirtus*)

From the 1,491 gene families expanded within *P. hirtus*, a total of 1,346 were annotated (Table S13). We observed the presence of several transferases, hydrolases, and ligases gene families, which were also identified in the other Elateroidea species. However, we recovered distinct unique or overrepresented molecular functions and biological processes in *P*.*hirtus* (Fig. S5F), such as multicellular organismal development, anatomical structure development and morphogenesis, reproduction, and multi-organism process (negative regulation of metabolic process). These terms are associated with the ontogenetic evolution of the organism, orchestrating the complex processes associated with embryogenic and growth development (Hill et al., 2010). The neoteny of the Phengodidae females and the winged male metamorphosis could be related to those gene families’ expansion associated with organismal ontogeny. The *superoxide dismutase* gene family was also expanded in Phengodidae, as well as *hypoxia-inducible factor 1-alpha*, which plays a role in the cellular response to O_2_ deprivation (Harrison et al., 2018). Several gene families displayed distinct orthologs in *P. hirtus*, such as *axin-1 like* (anterior development in Coleoptera; Fu et al., 2012), *chromobox protein homolog* (crucial in the establishment and maintenance of heterochromatin in larvae; Shoji et al., 2013), *doublesex- and mab-3-related transcription factor* (Sex determination mechanisms and sex differentiation; Rather and Dhandare., 2019), *forkhead box* (insulin signaling pathway and regulate physiological processes and juvenile hormone degradation; Zeng et al., 2017). Some other physiological mechanisms displayed overrepresented genes and we briefly discussed them below.

#### Extracorporeal digestion

The extracorporeal digestion is observed among distinct Insect families larvae, including the superfamily Elateroidea, in which this process was described in larvae of all bioluminescent species (Colepicolo-Neto et al. 1986; Bechara, 1988; Cohen et al., 1995). In the field, the Phengodidae larvae prey on millipede species (with equal or bigger corporeal size) or termites, introducing the digestive cocktail directly to the organism and, posteriorly, ingesting the partially digested fluid (Viviani and Bechara 1997). Colepicolo-Neto et al. (1986) showed a similar process in *P. termitilluminans* larva (Elateridae), identifying the presence of distinct proteases and carbohydrate hydrolases in the regurgitated fluid. Here, we identified the expansion of gene families associated with protease and carbohydrate hydrolases and extra-oral digestion in Arthropods, such as *chymotrypsin-1, chitinase, cathepsin Z, aminopeptidase N*, and *hyaluronidase*. These enzymes are also observed in other organisms, such as spiders (Fuzita et al., 2016; Walter et al., 2017). Some of these gene products are also present in the venom component of Heteropteran species and others (Fry et al., 2009; Walker et al., 2016). In particular, *hyaluronidase* seems to have an important role to increase the permeability of the tissue to the other toxins and digestive proteins (Walker et al., 2016).

#### Chemoreception

We identified olfactory, gustatory, odorant, and ionotropic receptors in expansion within the *P. hirtus* genome, suggesting the important role of intraspecific chemical communication in the Phengodidae. The chemosensory gene families expanded in the *P. hirtus* genome are associated with *gustatory and odorant receptor 22, olfactory receptor 2AG1, Glutamate receptor subunit 1*, and *Glutamate receptor ionotropic, kainate 2*. The chemosensory genes, in insects, are involved with mating, feeding, coordinating actions (e.g. attack, defense, escape), among others (Yuvaraj et al., 2018; Blomquist and Ginzel, 2021), being important physiological and ecological processes during the speciation process (mate isolation from its closely related species; Wu et al., 2019). The odorant and gustatory receptors are able to detect volatile chemicals, such as pheromones, which could be responsible for intraspecific communication, including mate. In Phengodidae, where the female is neotenic and the winged male displays a well-developed antenna (Costa et al., 1999), chemosensory communication is critical for sexual attraction. However, we did not observe the expansion of the protein binding gene families, the pheromones and odorant carriers, which indicates distinct evolutionary steps among the receptor and binding mechanisms in Phengodidae.

### 2.6. Gene families involved in *P. hirtus* bioluminescence

Based on previous molecular analysis, involving genomic and transcriptomic data, and biochemical studies (Niwa et al., 2006; Viviani et al., 2013; Oba et al., 2013; Kanie et al., 2016; Vongsangnak et al., 2016; Amaral et al., 2017a; 2017b; 2019b; Fallon et al., 2018; Zhang et al., 2020), we looked for specific described gene products that could be involved in the bioluminescence process in Elateroidea, mainly in *P. hirtus*. Here, we focused on luciferase evolution, bioluminescence emission control, antioxidant enzymes, and the luciferin biosynthesis pathway.

#### Luciferase evolution in Phengodidae

The number of AMP-forming enzymes observed in the Elateroidea species is abundant (ca. 15), including bioluminescent and non-bioluminescent species. In the last few years, transcriptomic and genomic data were able to recover the distinct isoforms of luciferase and luciferase-like enzymes and the comparison between their primary amino-acids sequence demonstrated the relationship among these isoforms and their evolution in Elateroidea. The presence of luciferase isozymes in the cephalic and lateral lanterns of Mastinocerini larvae was first shown by Viviani and Bechara (1993). Arnoldi et al. (2010), evaluating the relationship between the luciferase isoforms of Mastinocerini tribe species (Phengodidae), showed the presence of two luciferases, one in the cephalic lantern and one in the lateral body lanterns. These enzymes seem to be more closely related to the same lantern of different species than to the distinct lanterns of the same individual, suggesting an event of gene duplication and paralogy. Here, we identified 17 genes with similarities to luciferase and luciferase-like enzymes.

We were not able to completely assemble the genomic regions that contain the luciferase gene (only partial fragments). Thus, the raw reads were mapped against the Phengodidae luciferase and concatenated them. Using this strategy, we identified two genes that displayed 90% of similarity to Phengodidae luciferases, which were named PhLuc1 and PhLuc2. The similarity to the luciferase isoform described by Amaral et al. (2017) was ∼80%, which may suggest a non-specific gene assembly (possible chimera). The gene lengths for PhLuc1 is 2,019 bp (more similar to the lateral lanterns luciferase), while for PhLuc2 is 2,220 bp (more similar to cephalic lanterns luciferase), both genes comprising seven exons and six introns, which is a similar number of intergenic components observed for the luciferase of *Photinus pyralis* firefly (Fallon et al., 2018) and *Pyrophorus plagiophthalamus* click beetle (Elateridae; Feder and Vélez, 2009). The possible duplication event that originated both putative luciferases was followed by a dynamic structural genomic change, which altered the intron size, mainly the introns 1, 3, and 6, however, we did not observe evidence of intergenic combination. The average size of exons and introns were ∼230 pb and ∼100 pb, respectively.

#### Bioluminescence emission control and antioxidant enzymes

Recent studies using transcriptome and genomic data described possible gene products associated with the bioluminescent control in Elateridae, Lampyridae, and Phengodidae (Amaral et al., 2017a; 2017b; 2019b; Zhang et al., 2020). Trimmer et al. (2001) and Amaral et al. (2017b) suggested the importance of *nitric oxide synthase* and *octopamine/dopamine receptors* to firefly flash control, mainly in the adult stage, in which the flash pattern and duration are fundamental for intraspecific communication. However, these gene products were less abundant in Elateridae and Phengodidae, which is consistent with the lower degree of neural control and more continuous glow pattern in these species (Amaral et al., 2017a; 2019b). Here, only a gene of *nitric oxide synthase* and *dopamine/octopamine receptor* were found. In Lampyridae genomes, only a copy of the nitric *oxide synthase* gene was also identified, suggesting a unique gene associated with the control of available oxygen concentration inside the photocytes (Trimmer et al., 2001). The number of *dopamine/octopamine receptors* was between 8 to 10 copies, much higher than observed in phengodids, consistently with the need for flash control in the adult stage of fireflies.

The presence of *catalase* and *superoxide dismutase* in photophores of Elateroidea was previously described by Barros and Bechara (1998) and was associated with detoxification reactive oxygen species (ROS) inside the cells. We already described the abundance of these gene products in the Elateroidea lanterns using a transcriptome dataset and, in this study, we identified two genes of *catalase* and three of *superoxide dismutase Cu-Zn* (SOD). The SOD enzyme catalyzes the dismutation of the superoxide radicals to molecular oxygen and H_2_O_2_, while the catalase dismutates the H_2_O_2_ into O_2_, avoiding harmful damage to the photocytes (Tainer et al., 1983; Cox et al., 2011).

#### Luciferin biosynthesis

In *P. hirtus* genome, several gene products that were previously potentially associated with luciferin biosynthesis in luminescent Elateroidea were found (Niwa et al., 2006; Viviani et al., 2013; Vongsangnak et al., 2016; Amaral et al., 2017a, 2017b, 2019b; Zhang et al., 2020). Among them, the *adenosylhomocysteinase* and *cysteine sulfinic acid decarboxylase* are associated with the conversion of homocysteine to cysteine, which may spontaneously react with *p*-benzoquinone to generate luciferin (Kanie et al., 2016). The gene products involved with tyrosine metabolism (*tyrosine aminotransferase, tyrosine hydroxylase, 4-hydroxyphenylpiruvate dioxygenase*, and *homogentisate 1,2-dioxygenase*) and the cascade reaction of the L-DOPA pathway (*dopamine/octopamine receptor, dopamine N-acetyltransferase, sodium-dependent dopamine transporter*, and *phenoloxidase* and *phenoloxidase activating factor*) were also observed, as well as a *luciferin-regenerating enzyme* gene (LRE), which was suggested to recycle the oxyluciferin to an intermediary compound of *L-Luciferin* (*2-cyano-6-hydroxybenzothiazole*), and a gene of luciferin *sulfotransferase*, which converts luciferin to a stable storage compound, *sulfoluciferin* (Fallon et al., 2016). However, we did not recover any *Acyl-CoA thioesterase* (ACT) gene which could be involved in the conversion of *L-luciferin* to *D-luciferin* (Niwa et al., 2006), whereas we observed the expansion of this gene family among Lampyridae. Rather, we identified two genes of *palmitoyl-protein thioesterase* (a specific group withn ACT), which is a lysosomal enzyme that removes fatty acyl groups from cysteine residues (Glaser et al., 2003), similar to the function of the ACT genes in peroxisomes (Lousa et al., 2013).

These results suggest a quite similar luciferin biosynthetic pathway within Elateroidea bioluminescent species, however, some steps or genes involved in the cascade reactions may have evolved differently during the diversification of the Elateroidea families. Thus, the cellular and metabolic processes may have favored one enzyme over another, such as the *palmitoyl-protein thioesterase* instead of the *Acyl-CoA thioesterase*.

## 3. Concluding remarks

We combined short and long reads generated with Illumina sequencing to generate the first draft genome of the South-American *Phrixothrix hirtus* railroad worm, the first one of the Phengodidae family. This is the largest genome observed in the Elateroidea superfamily, with more than 60% of its size populated by TE, including the presence of several retrotransposable elements, such as LINE, MOS1, and PiggyBac. The *P. hirtus* genome displays unique gene families related to anatomical development and morphogenesis, consistently with the neoteny and strong sexual dimorphic development in this species.

The comparison of gene families orthogroups among Elateriformia showed quite similar molecular functions and biological processes shared among all species, which are enriched by catalytic activity and binding enzymes, but did not show clear differences in the presence or absence of gene families associated with the bioluminescence among species/families. A very similar pattern of genes putatively involved in luciferin biosynthetic pathway within Elateroidea bioluminescent species was observed, with the exception of the esterase involved in the conversion of *L-* to *D-luciferin*, which is a *palmitoyl-thioesterase* in *P. hirtus* instead of an *acyl-esterase* as found in Lampyridae. These results suggest that genes involved in bioluminescence production may have evolved before the divergence within the Elateriformia, whereas the spatial and temporal transcriptional and expression control levels may have evolved later determining the bioluminescence anatomical pattern distribution and control rather than the genomic features.

## 4. Material and methods

### 4.1. Sampling, DNA extraction, and libraries construction

One larval individual of *P. hirtus* was manually collected at “Fazenda Santana” (Souza, Campinas/SP, Brazil) and identified by Prof. Dr. Vadim Viviani. The sample was stored at -80°C until the DNA extraction. The genomic DNA was extracted from whole larvae using DNeasy blood & tissue kit (Qiagen, USA), according to the instructions of the manufacturer. The DNA quantity and quality were measured using NanoVue (GE HealthCare) and Qubit 3.0 fluorometer (ThermoFisher, USA), and the integrity was checked in agarose gel. Two genomic DNA libraries were prepared: i. using the TruSeq DNA PCR-free library prep kit with fragments of 150 bp and ii.) Nextera Mate Pair library prep with fragments of 2,000 bp (Illumina, USA). The short-read paired-end libraries were sequenced in two independent lanes and the mate-pair library in one lane using the Illumina HiSeq4000 platform (Illumina, USA). The library’s construction and sequencing were performed by Hokkaido System Science Co. (Sapporo, Hokkaido, Japan). The raw read data and final genome assembly of *P. hirtus* are deposited and available at the BioProject PRJNA741915.

### 4.2. Pre-processing data, *de novo* genome assembly, and annotation

The reads obtained by the DNA-Seq were checked by FastQC v0.11.6 software (http://www.bioinformatics.babraham.ac.uk/projects/fastqc/), and adaptors and low-quality reads (Phred Q ≤ 30) were removed using FASTX-TOOLKIT v0.0.14 (Pearson et al., 1997) for the paired-end library and NxTrim (O’Connell et al., 2015) for the mate-pair library. We used DeconSeq v.0.4.3 (Schmieder and Edwards, 2011) software and the RefSeq database (bacteria and viruses) (accessed in April 2018) to remove any microbial contamination within the raw data sequencing. After these filtering processes, we proceed to downstream genomic assembly.

We estimated the best k-mer length for the *P. hirtus* genome assembly using KmerGenie 1.705 (Chikhi and Medvedev, 2014), which was k=81 (data not shown). The genome coverage, size heterozygosity were estimated with *estimate_genome_size*.*pl* script (available at https://github.com/josephryan/estimate_genome_size.pl), Jellyfish2 v.2.2.3 (Marcais and Kingsford, 2011) (parameters: *count -t 8 -C -m 21 --min-quality=20 --quality-start=33*), and GenomeScope2 (Vurture et al., 2017; available at http://qb.cshl.edu/genomescope/genomescope2.0/). We used the gatb-minia-pipeline (available at https://github.com/GATB/gatb-minia-pipeline) for the *de novo* assembly using default settings and fixing k-mer size in 81. The scaffolded statistics were evaluated using QUAST 5.0.0 (Gurevich et al., 2013). For the mitochondrial genome assembly, we used GetOrganelles software (Jin et al., 2020), with the default setting, and MITOS Web Server (Bernt et al, 2013) for the annotation.

The gene prediction was performed using two-round training in MAKER software with the default settings, combined with transcriptome-based nucleotide and amino acid sequences generated by Amaral et al. (2017a) to improve the prediction. Repetitive DNA elements were identified and annotated using RepeatModeler v.1.0.8 (see http://www.repeatmasker.org/RepeatModeler), RepeatMasker v.4.0.9 (see http://www.repeatmasker.org/). The annotation of the coding regions was conducted using the tool BLASTX against the Swissprot database (retrieved on 05/2020). The GO terms were plotted and visualized using WEGO2.0 (Ye et al., 2018). The completeness of the genes was estimated using BUSCO v.3.0.2 software (Simão et al., 2015).

### 4.3. Orthologous protein clustering, gene family evolution, and phylogenetic analysis

Available genomic data from seven Elateriformia species (details in Table S14) were utilized to perform comparative genomic analyses. The orthogroups identification and the expanded and contracted gene families were identified with OrthoFinder v.2.0.9 (Emms and Kelly, 2015) and CAFÉ v.4.2.1, using birth and death rate (Han et al., 2013), respectively. The ultrametric tree used to determine expansion and contraction family size was performed using r8s v.1.81 software (Sanderson, 2003). The phylogenetic relationship topology was estimated using 359 conserved single-copy orthologs in IQtree2 (Minh et al., 2020) in 1,000 ultrafast bootstraps.

### 4.4. Luciferase genes identification

To identify the putative luciferase gene length, the raw reads were mapped against the luciferase of Phengodidae species available at public databases, using the bowtie2 v.2.4.3 tool (Langmead and Salzberg, 2012). We processed the reads using Samtools v.1.9 (Li et al., 2009) and concatenated them using CAP3 v.10.2011 (Huang and Madan, 1999) software.

## Acknowledgments

We thank the Fundação de Amparo à Pesquisa do Estado de São Paulo (FAPESP grant 2010/05426-8 to VV; FAPESP n. 2017/207340 and 2014/20176-9 to DTA) for Financial Support.

## Author contributions

VV, YO, YM, and DTA conceived the idea; DTA and IASB led both the analyses and manuscript writing; VV, YO, YM, and CR collaborated in manuscript writing. All authors contributed to the intellectual development of the paper, made multiple revisions, and approved the final draft.

## Competing interest

The authors declare no competing interest.

## Notes

### Competing Interest Statement

The authors have declared no competing interest.

