## Supplementary material for "Draft genome of the Brazilian railroad worm *Phrixothrix hirtus* E.Olivier (Phengodidae: Coleoptera)": SM1

**^1^** Department of Biology. Center for Human and Biological Sciences. Universidade Federal de São Carlos (UFSCar). Sorocaba, Brazil.

^2^ Graduate Program in Comparative Biology. Faculty of Philosophy, Sciences, and Languages of Ribeirão Preto. Universidade de São Paulo (USP). Ribeirão Preto, Brazil.

^3^ Bioproduction Research Institute, National Institute of Advanced Industrial Science and Technology (AIST), Sapporo, Japan.

^4^ Department of Ecology and Evolutionary Biology. Universidade Federal de São Paulo (UNIFESP). Diadema, São Paulo, Brazil.

^5^ Department of Computational Science, Universidade Federal de São Carlos (UFSCar). São Carlos, Brazil.

^6^ Biomedical Research Institute, AIST, Ikeda-Osaka, Japan.

^7^ Osaka Institute of Technology, OIT, Osaka, Japan.

^8^ Graduate Program of Evolutive Genetics and Molecular Biology, Federal University of São Carlos (UFSCar), São Carlos, Brazil.

^9^ Graduate Program of Biotechnology and Environmental Monitoring, Federal University of São Carlos (UFSCar), Sorocaba, Brazil.

*Corresponding author: Rodovia João Leme dos Santos, Km 110, SP264. ZIP 18052-780, Sorocaba, Brazil. Phone: +55-015-3229-5983, Fax: + 55-015-3229-5983.; ^#^

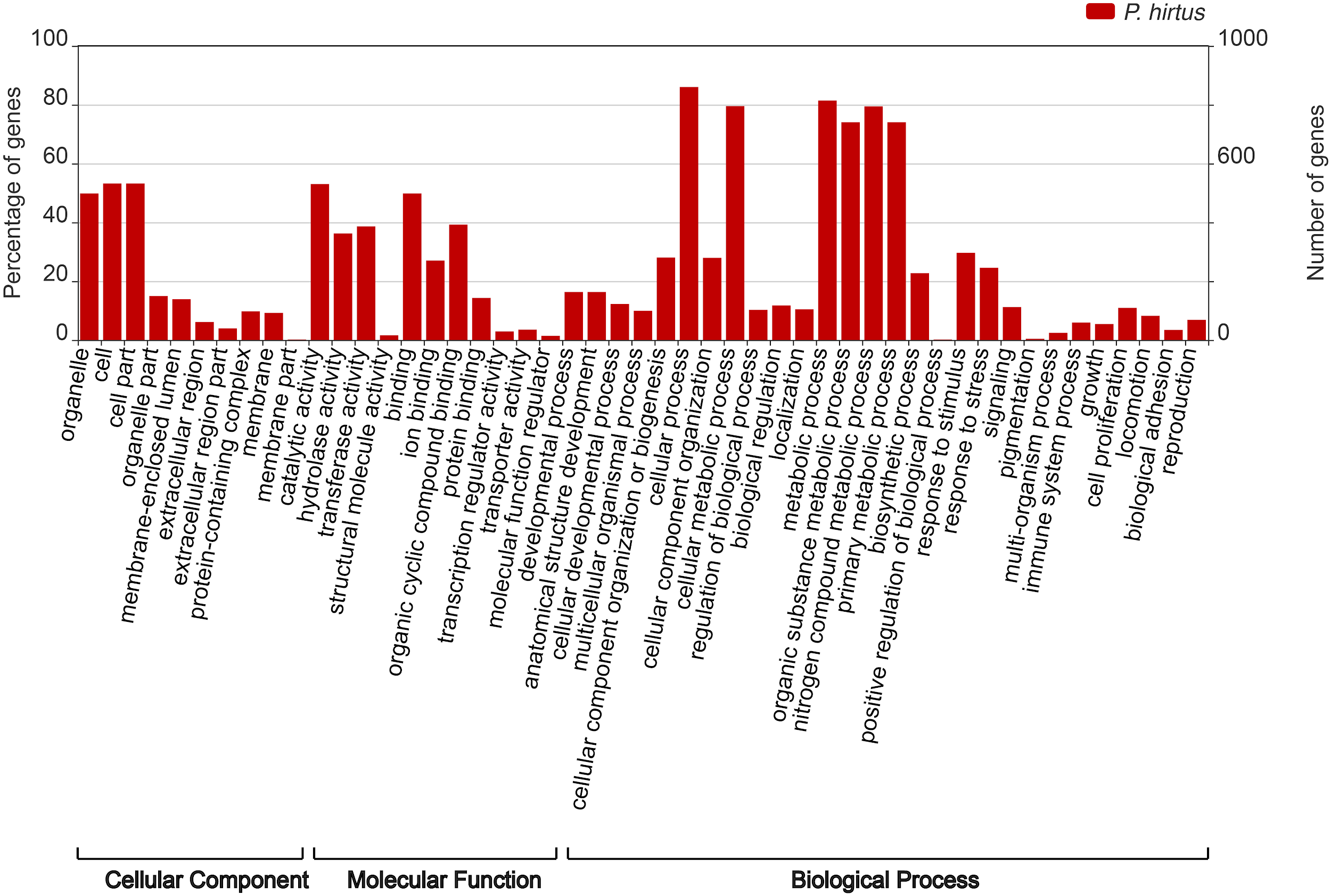

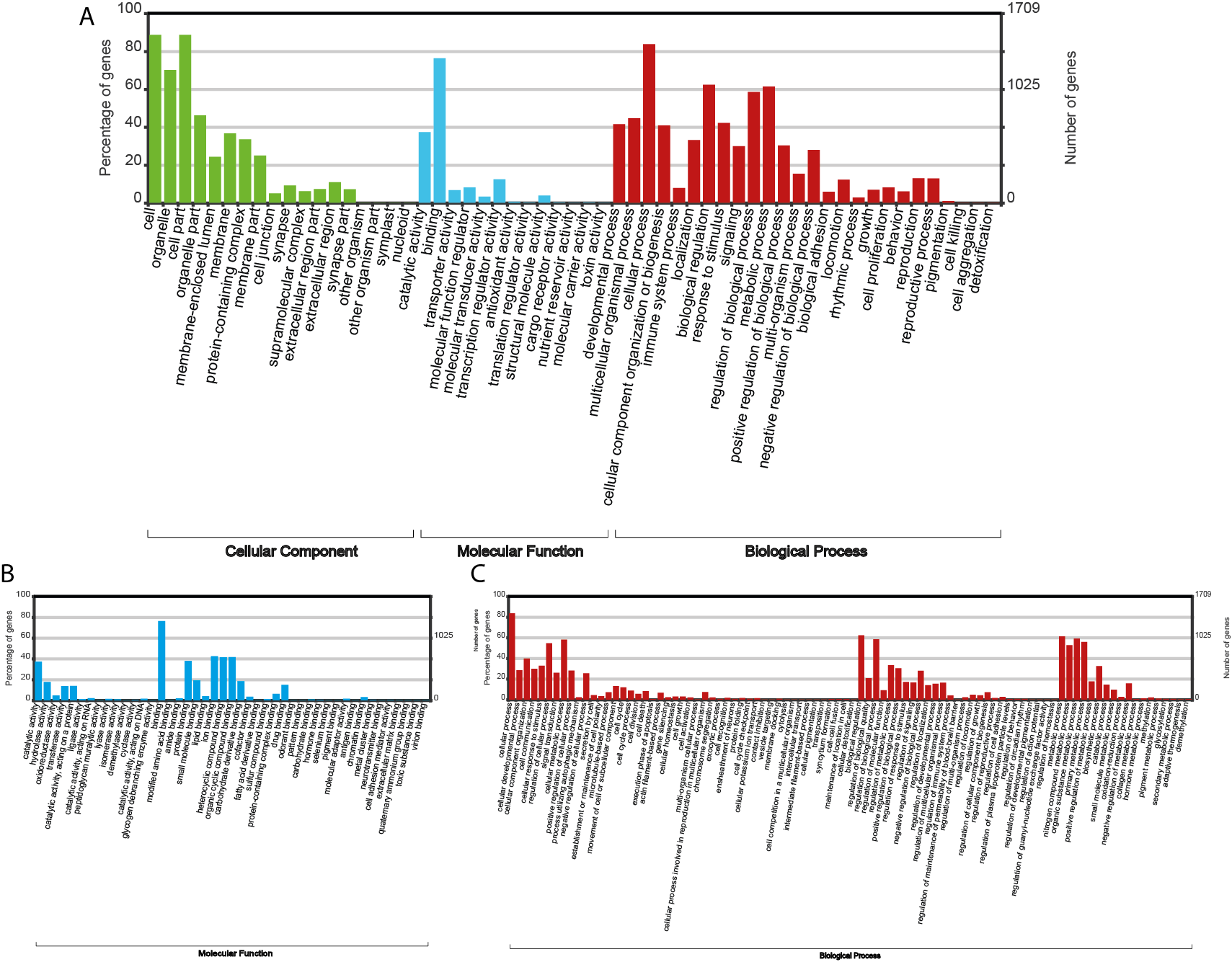

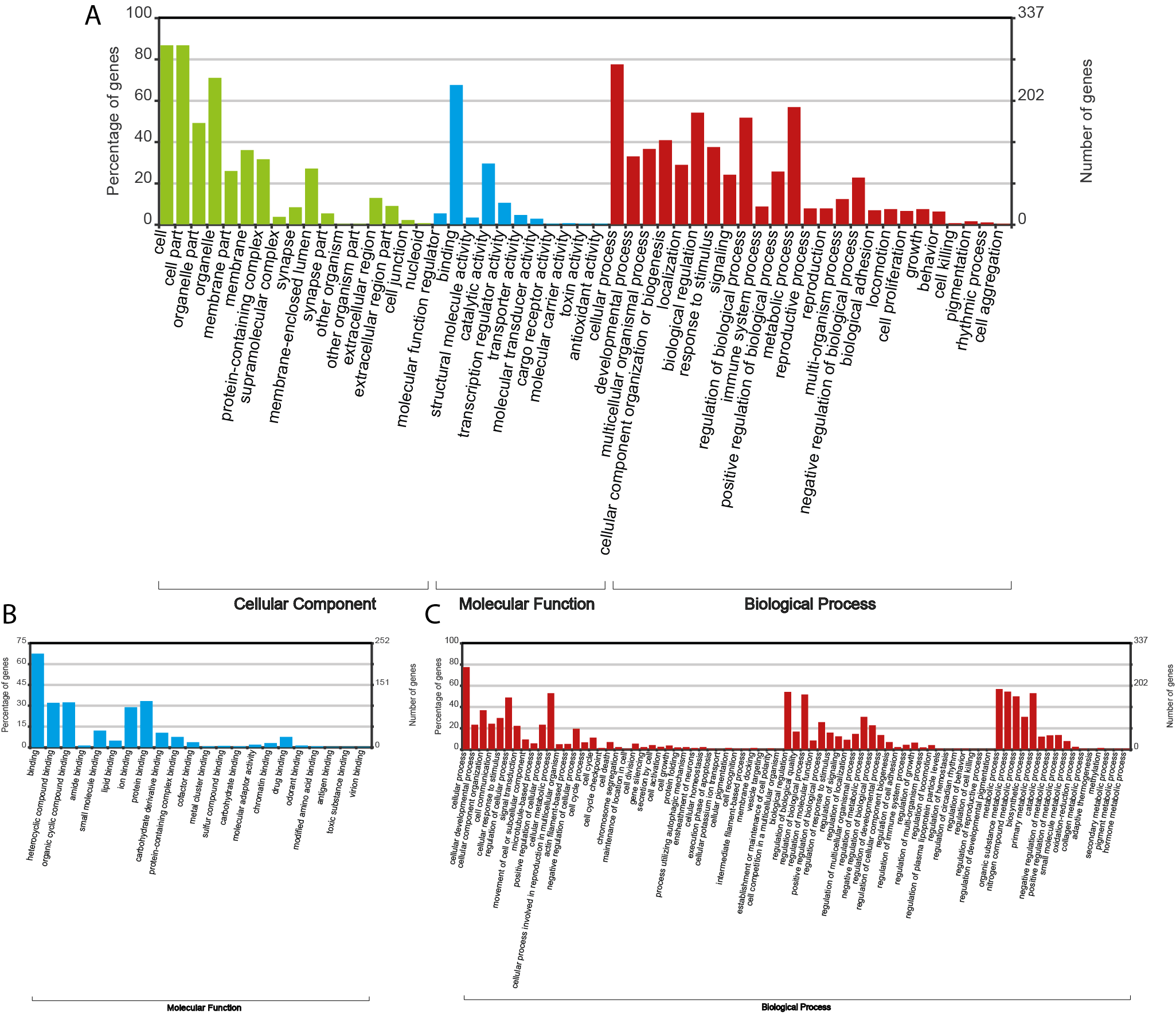

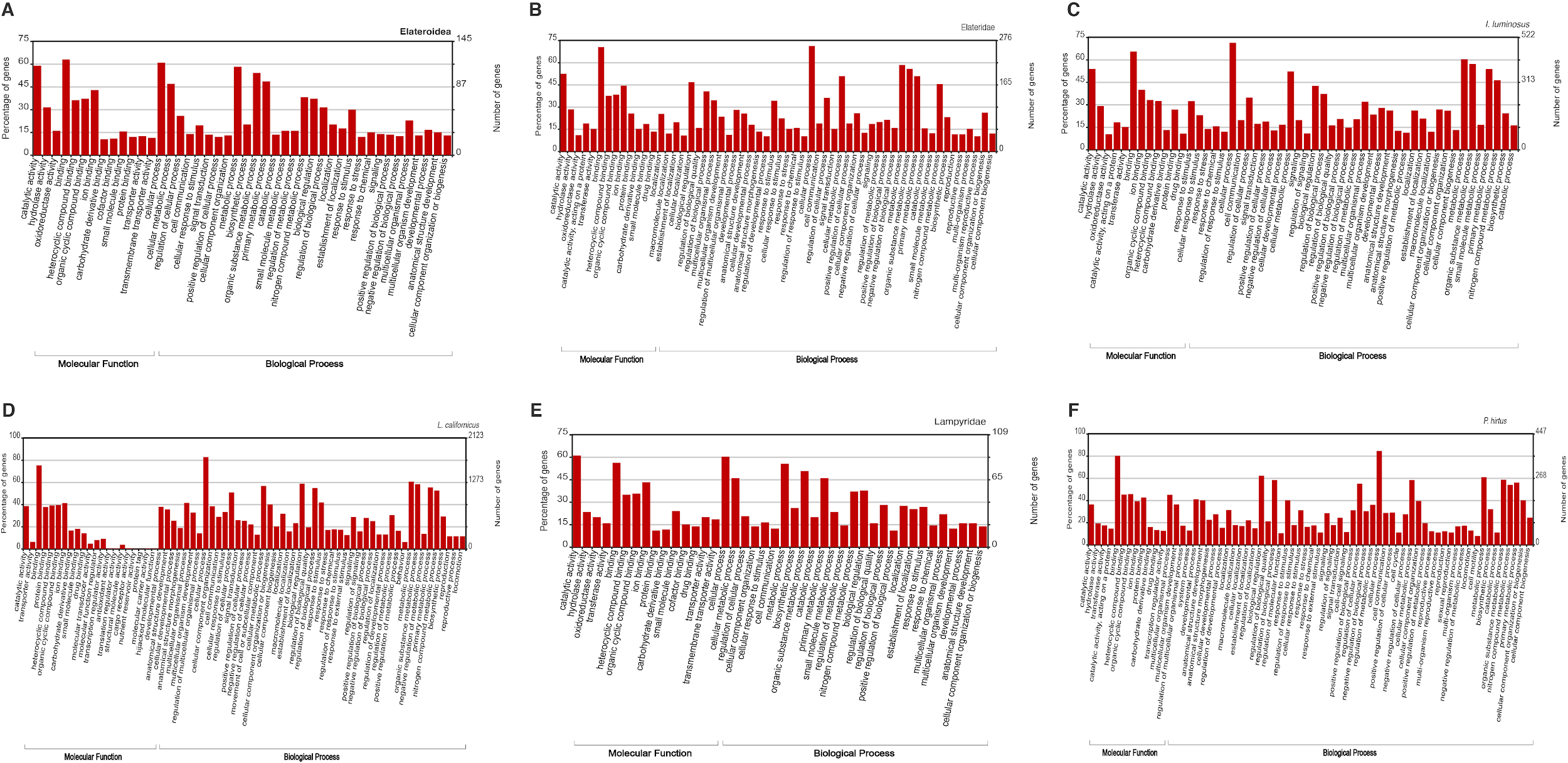
